# Kinetochore components are perinuclear and form condensates in *C. elegans* germline

**DOI:** 10.64898/2025.12.16.694141

**Authors:** Hannah L. Hertz, Ian F. Price, Wen Tang

## Abstract

The kinetochore is a multiprotein complex that assembles on centromeric chromatin and serves as the attachment site for spindle microtubules, thereby ensuring accurate chromosome segregation during cell division. Here we present evidence that kinetochore components, such as HCP-1, form distinct cytoplasmic foci in the *C. elegans* germline. Prior to cell division, HCP-1 localizes to the nuclear envelope of germ nuclei and forms granules that are distinct from germ granules. Upon mitotic entry, perinuclear HCP-1 is imported into the nucleus to facilitate chromosome segregation. We further show that HCP-1 associates with nuclear pore proteins (NPPs). Loss of NPP-14 disperses perinuclear HCP-1, impairs its nuclear import, and leads to defects in germ cell proliferation and embryonic spindle assembly. Together, these findings provide new insights into the spatial regulation of kinetochore components and their roles in nuclear organization.

## Introduction

The kinetochore complex is a conserved, multi-protein complex essential for the accurate segregation of chromosomes during cell division (Cheeseman, 2014; Cheeseman and Desai, 2008; McAinsh and Marston, 2022). The complex assembles on centromeric chromatin and mediates the attachment of chromosomes to spindle microtubules and facilitates chromosome movement during both mitosis and meiosis (Foley and Kapoor, 2013; Musacchio and Desai, 2017). The kinetochore structure is not only responsible for physically linking chromosomes to spindle microtubules, but also for generating the forces necessary to align and segregate chromosomes accurately. Defects in kinetochore assembly can lead to chromosome mis-segregation and aneuploidy, which contributes to developmental disorders, infertility and cancer (Santaguida and Amon, 2015).

In contrast to monocentric organisms with point-shaped kinetochores, the nematode *C. elegans* has holocentric chromosomes in which kinetochores assemble along the entire length of the chromosome (Albertson and Thomson, 1982). Despite this structural difference, *C. elegans* expresses many conserved kinetochore components and share similar features in the molecular composition, assembly, and function of kinetochores with other organisms (McIntosh et al., 2013; Pintard and Bowerman, 2019). Broadly, there are inner and outer kinetochore proteins. The inner kinetochore proteins, including histone H3 variant known as HCP-3 in *C. elegans* or CENP-A in vertebrates, associate with centromeric DNA (Buchwitz et al., 1999). The outer proteins, such as HCP-1 and HCP-2 in *C. elegans* or CENP-F in vertebrates, connect kinetochores with microtubules (Cheeseman et al., 2005; Moore et al., 1999). Therefore, the *C. elegans* kinetochore serves as an excellent model for understanding how chromosome-microtubule attachments are spatially and temporally regulated and provides insights into conserved mechanisms of chromosome segregation across species.

Germ granules were characterized as perinuclear and membrane-less biomolecular condensates that are essential for germline development and fertility in diverse animals (Brangwynne et al., 2009; Mahowald, 1968; Strome and Wood, 1982). In *C. elegans*, several distinct but interconnected types of germ granules have been described, including P granules, Mutator foci, E granules, and Z granules (Chen et al., 2024; Phillips et al., 2012; Wan et al., 2018). Collectively, these germ granules control small RNA synthesis, mediate gene silencing, participate in epigenetic inheritance, and are required for maintaining germline fate (Phillips and Updike, 2022). Among those, P granules were the first identified and best characterized (Strome and Wood, 1982). P granules are positioned at the cytoplasmic face of nuclear pores in germ cells (Pitt et al., 2000; Updike et al., 2014). Nuclear pores consist of over 30 different proteins referred to nucleoporins (Nups) in humans or nuclear pore proteins (NPPs) in *C. elegans* (Strambio-De-Castillia et al., 2010). Notably, NPPs form clusters the nuclear envelope of germ nuclei (Pitt et al., 2000; Sheth et al., 2010; Shi et al., 2025; Thomas et al., 2025). In addition, a subset of Nups/NPPs form nucleoporin foci in developing oocytes (Patterson et al., 2011; Pitt et al., 2000; Thomas et al., 2023). However, the molecular function and composition of nucleoporin foci have not been fully elucidated.

Using a proximity-based approach, we recently defined the protein composition of P granules (Hertz et al., 2022; Price et al., 2021). As expected, we identified canonical P granule components and several NPPs (Hertz et al., 2022; Price et al., 2021). Unexpectedly, we also detected multiple kinetochore proteins including HCP-1. In this study, we show that HCP-1 assembles into perinuclear granules at germ nuclei and cytoplasmic condensates in oocytes. Perinuclear HCP-1 translocates into nuclei during mitosis, where it contributes to kinetochore formation. Loss of NPP-14 leads to the dispersal of perinuclear HCP-1 and impairs its nuclear reimport. Together, these findings uncover a previously unrecognized layer of spatial regulation of kinetochore components and suggest a functional link between nuclear pore proteins and kinetochore dynamics in the germline.

## Results

### HCP-1 forms cytoplasmic foci that colocalize with nucleoporins

Our previous proximity labeling studies defined the proteome of *C. elegans* P granules (Hertz et al., 2022; Price et al., 2021). In these experiments, known P granule proteins such as GLH-1 were fused with a promiscuous biotin ligase that biotinylates proximate proteins. Biotinylated proteins were enriched in adult worms and further identified using mass spectrometry (Hertz et al., 2022; Price et al., 2021). Protein-protein interaction network analysis of this dataset revealed at least four clusters including a germ granule cluster, a NPP cluster, and a kinetochore cluster (Figure S1A). Identification of P granule and nuclear pore components was expected, as P granules situate at the cytoplasmic face of nuclear pore complexes (Pitt et al., 2000; Sheth et al., 2010; Thomas et al., 2025). However, the presence of a kinetochore cluster was unexpected (Figure S1A), as kinetochores are believed to associate with chromosomes hence localize to the nucleus. Among the kinetochore components, HCP-1—a homolog of vertebrate CENP-F (Cheeseman et al., 2005; Moore et al., 1999)—showed the strongest enrichment from our proximity labeling experiments (Figure S1B). By contrast, HCP-2, the HCP-1 paralog, was not significantly enriched (Figures S1A and S1B).

To validate this surprising finding, we examined the localization of endogenously GFP-tagged HCP-1 in both embryos and adult gonads (Edwards et al., 2018; Macaisne et al., 2023). The adult *C. elegans* gonad is a syncytial tissue in which germ cells undergo mitotic proliferation at the distal end and then enter meiotic pachytene to form oocytes. Oocytes complete meiosis after fertilization, and the resulting zygote undergoes rapid cell divisions to initiate embryonic development (Figure S1C) (Pazdernik and Schedl, 2013). In line with its role in kinetochore function (Cheeseman et al., 2005; Edwards et al., 2018), GFP::HCP-1 was robustly expressed in nuclei and formed kinetochore structures in dividing embryonic cells (Figure S1D). In the adult germline, GFP::HCP-1 was also expressed, with high levels observed in the mitotic zone and developing oocytes, and lower levels in the meiotic pachytene region (Figure 1A). In contrast to its nuclear localization in embryos (Figure S1D), GFP::HCP-1 primarily formed perinuclear foci in meiotic germ cells (Figures 1A, 1B and S1C), and perinuclear and cytoplasmic foci in oocytes (Figures 1A, 1C and S1C).

**Figure 1.**
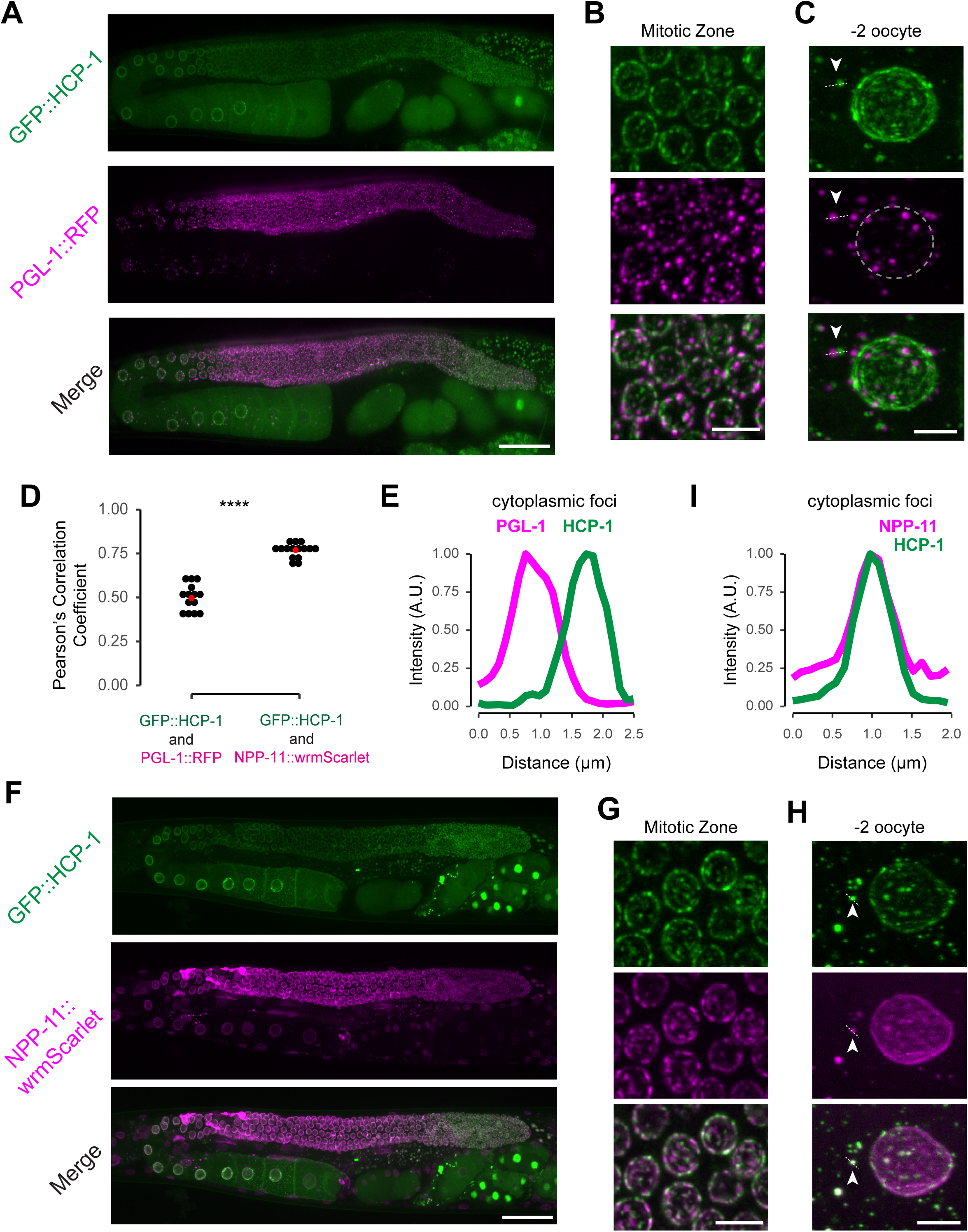
HCP-1 forms cytoplasmic foci that colocalize with nucleoporin proteins. **A.** Representative maximum intensity projection (40x objective) of GFP::HCP-1 and PGL-1::RFP in the whole germline of a day 2 adult. Scale bar: 40 µm. **B.** Maximum intensity projection (60x objective) of germ nuclei at the mitotic zone for the strain in A. Scale bar: 5 µm. **C.** Maximum intensity projection (60x objective) of a -2 oocyte for the strain in A. Dashed circle outlines the nuclear periphery. Arrow indicates the cytoplasmic focus analyzed in E. Scale bar: 5 μm. **D.** Quantification of correlation by the Pearson’s Correlation for the indicated proteins. Each black dot represents the Pearson’s R-value for the green vs. red channel of a single nucleus. Red dot indicates the mean. Statistics determined using Wilcoxon rank-sum test and **** represents p<0.0001. **E.** Intensity profile of GFP::HCP-1 (green) or PGL-1::RFP (magenta) cytoplasmic focus along the dashed line in C. **F.** Representative maximum intensity projection (40x objective) of GFP::HCP-1 and NPP-11::wrmScarlet in the whole germline of a day 2 adult. Scale bar: 40 µm. **G.** Maximum intensity projection (60x objective) of germ nuclei at the mitotic zone for the strain in F. Scale bar: 5 µm. **H.** Maximum intensity projection (60x objective) of a -2 oocyte for the strain in F. Dashed circle outlines the nuclear periphery. Arrow indicates the cytoplasmic focus analyzed in I. Scale bar: 5 μm. **I.** Intensity profile of GFP::HCP-1 (green) or NPP-11::wrmScarlet (magenta) cytoplasmic focus along the dashed line in panel H.

To further determine the subcellular localization of HCP-1, we co-expressed PGL-1::RFP, an established P granule marker (Kawasaki et al., 1998; Wan et al., 2018). We found that GFP::HCP-1 formed foci that only partially overlapped with PGL-1::RFP (Figures 1A-1C). The relationship between GFP::HCP-1 and PGL-1::RFP foci was measured as Pearson’s correlation coefficient, where 1 indicates perfect correlation, 0 indicates no correlation, and –1 indicates perfect anti-correlation. This analysis revealed only modest colocalization between HCP-1 and PGL-1 (mean Pearson’s correlation coefficient = 0.50) (Figure 1D). This observation was further supported by visual inspection of individual GFP::HCP-1 and PGL-1::RFP cytoplasmic foci in oocytes. Specifically, quantification of fluorescence intensity confirmed that HCP-1 and PGL-1 form overlapping but distinct structures (Figures 1C and 1E).

The expression pattern of GFP::HCP-1 resembled that of nuclear pore proteins (NPPs), which are typically enriched at the nuclear envelope. Notably, a subset of NPPs including NPP-11 and NPP-14 form nucleoporin foci in growing oocytes (Patterson et al., 2011; Pitt et al., 2000; Thomas et al., 2023). To test whether HCP-1 associates with these structures, we co-expressed GFP::HCP-1 and NPP-11 fused with wrmScarlet (Thomas et al., 2023). GFP::HCP-1 and NPP-11::wrmScarlet displayed strong colocalization at the nuclear periphery of miotic and meiotic germ cells (Figures 1F-1H). Notably, the mean Pearson’s correlation coefficient for GFP::HCP-1 and NPP-11::wrmScarlet was 0.77, significantly higher than that observed between GFP::HCP-1 and PGL-1::RFP (Figure 1D). Quantification of the cytoplasmic foci in oocytes confirmed substantial colocalization between GFP::HCP-1 and NPP-11::wrmScarlet (Figure 1I). Similarly, GFP::HCP-1 colocalized with another NPP (NPP-14::wrmScarlet) in both mitotic germ nuclei as well as oocytes (Figures S2A-D). Together, our findings suggest that HCP-1, a kinetochore component, forms both perinuclear and cytoplasmic foci with NPPs in the *C. elegans* germline.

### Distinct properties of perinuclear and cytoplasmic HCP-1 foci

Next, we asked whether HCP-1 foci exhibit properties of liquid-like droplets. At least two key features define a liquid-like condensates: (1) a 2-dimensional rounded or 3-dimensional spherical morphology due to the surface tension, and (2) dynamic molecular exchange with the surrounding environment (Brangwynne et al., 2009; Patel et al., 2015). We examined these two features in perinuclear HCP-1 foci located in the mitotic germ nuclei and cytoplasmic HCP-1 foci present in developing oocytes. To assess shape, we quantified the roundness of individual foci, where a value of 1 represents a perfect circle (Figure 2A). We found that cytoplasmic HCP-1 foci were significantly more circular than their perinuclear counterparts (Figure 2A).

**Figure 2.**
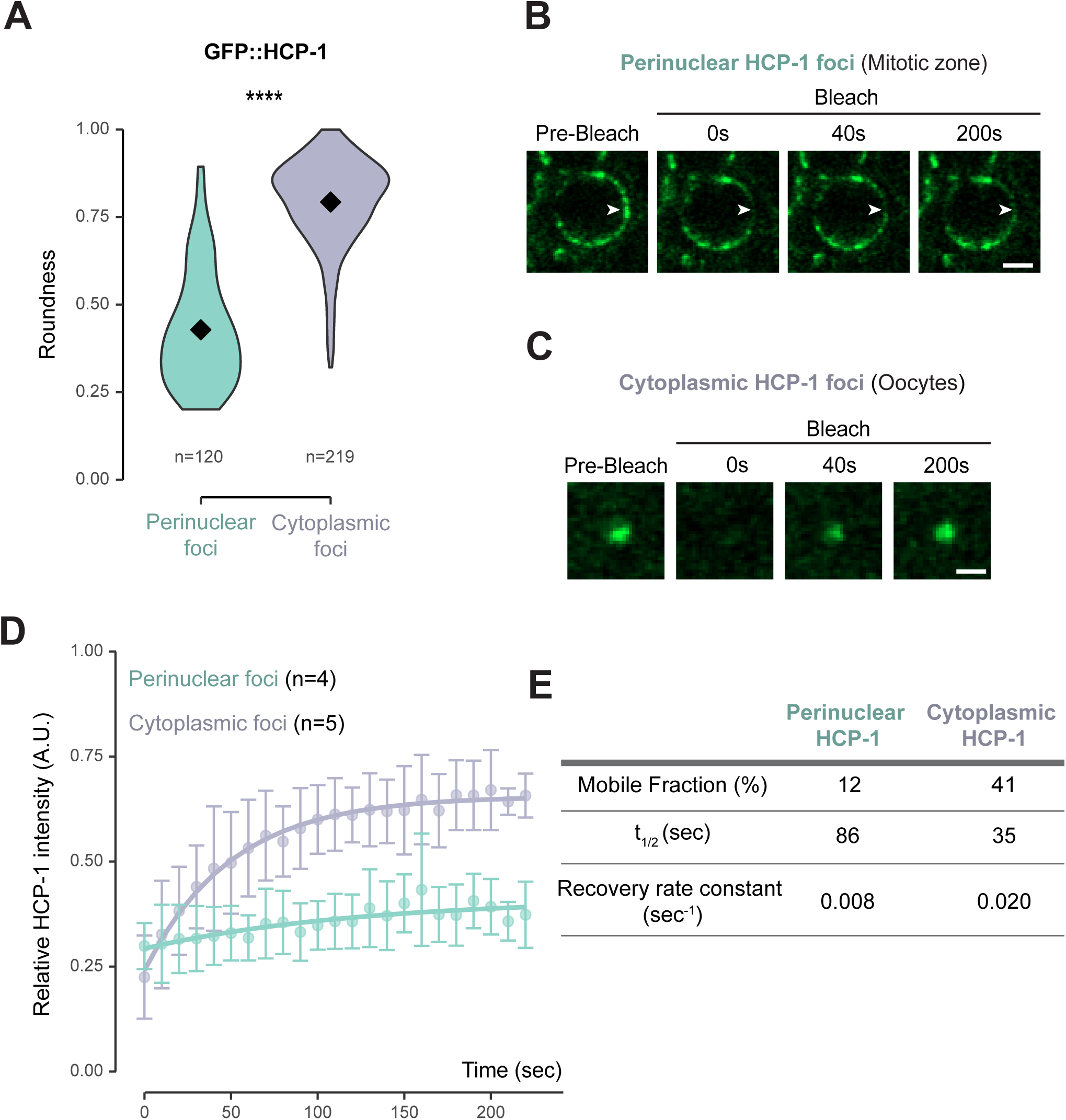
Distinct properties of perinuclear and cytoplasmic HCP-1 foci. **A.** Violin plot of roundness measurement for perinuclear (teal) or cytoplasmic (purple) GFP::HCP-1 foci. Diamond represents the median. *n* refers to each focus analyzed. Statistics determined using Wilcoxon rank-sum test and **** represents p<0.0001. **B.** Single-plane timelapse (63x objective) of a nucleus at the mitotic zone showing GFP::HCP-1 fluorescence before (pre-bleach) and after photobleaching at time zero, at the indicated timepoints (seconds). Arrow indicates the bleached perinuclear focus. Scale bar: 2 µm. **C.** Single-plane timelapse (63x objective) of a cytoplasmic focus in an oocyte showing GFP::HCP-1 fluorescence before (pre-bleach) and after photobleaching at time zero, at the indicated timepoints (seconds). Scale bar: 1 µm. **D.** Graph showing fluorescence recovery after photobleaching (FRAP) for GFP::HCP-1 perinuclear foci (teal, n=4) or cytoplasmic foci (purple, n=5). Intensity was measured from single-plane images every 10 seconds (sec) before and after photobleaching. Relative HCP-1 intensity at each timepoint was normalized to the mean HCP-1 foci intensity prior to photobleaching. Bleaching occurred at timepoint 0. Error bars represent the mean +/- sd. Teal or purple curves represent the first-order model of HCP-1 recovery for perinuclear or cytoplasmic foci, respectively. **E.** Table showing recovery of perinuclear HCP-1 foci compared to cytoplasmic HCP-1 foci after photobleaching. The mobile fraction, t_1/2_, and recovery rate constant was derived from the models in D.

Furthermore, we assessed the dynamics of HCP-1 foci using fluorescence recovery after photobleaching (FRAP). In brief, GFP::HCP-1 foci were photobleached and fluorescence recovery was monitored over 220 seconds (Figures 2B and 2C), and recovery curves were fitted to a single exponential model (Figure 2D). We found cytoplasmic GFP::HCP-1 foci displayed faster and more complete recovery than perinuclear foci (Figures 2B–2D). Specifically, the half-time of recovery (t_1/2_) was 35 seconds for cytoplasmic HCP-1 and 86 seconds for perinuclear HCP-1 (Figure 2E). The mobile fraction was also higher for cytoplasmic GFP::HCP-1 (41%) compared to perinuclear GFP::HCP-1 (12%) (Figure 2E). Together, these data revealed distinct properties of perinuclear and cytoplasmic HCP-1 foci with cytoplasmic foci in growing oocytes being more dynamic and liquid-like.

### Formation of HCP-1 foci and P granules are independent

To investigate whether HCP-1 foci and P granules are interdependent, we used RNA interference (RNAi) to deplete key components of P granules in animals expressing both PGL-1::RFP and GFP::HCP-1 (Fire et al., 1998; Kamath et al., 2003), Specifically, we targeted the Vasa homologs GLH-1 and GLH-2, as well as the Argonaute protein CSR-1. Depletion of GLH-1/2 disrupted perinuclear P granules and led to the formation of large cytoplasmic granules (Figures 3A and 3B), consistent with previous reports (Cipriani et al., 2021; Price et al., 2021; Price et al., 2023; Updike and Strome, 2009). Loss of CSR-1 resulted in enlarged and irregular P granules (Figures 3A and 3B) (Claycomb et al., 2009; Updike and Strome, 2009). In contrast, depletion of either GLH-1/2 or CSR-1 did not appear to strongly affect the formation or localization of GFP::HCP-1 foci (Figures 3A and 3B).

**Figure 3.**
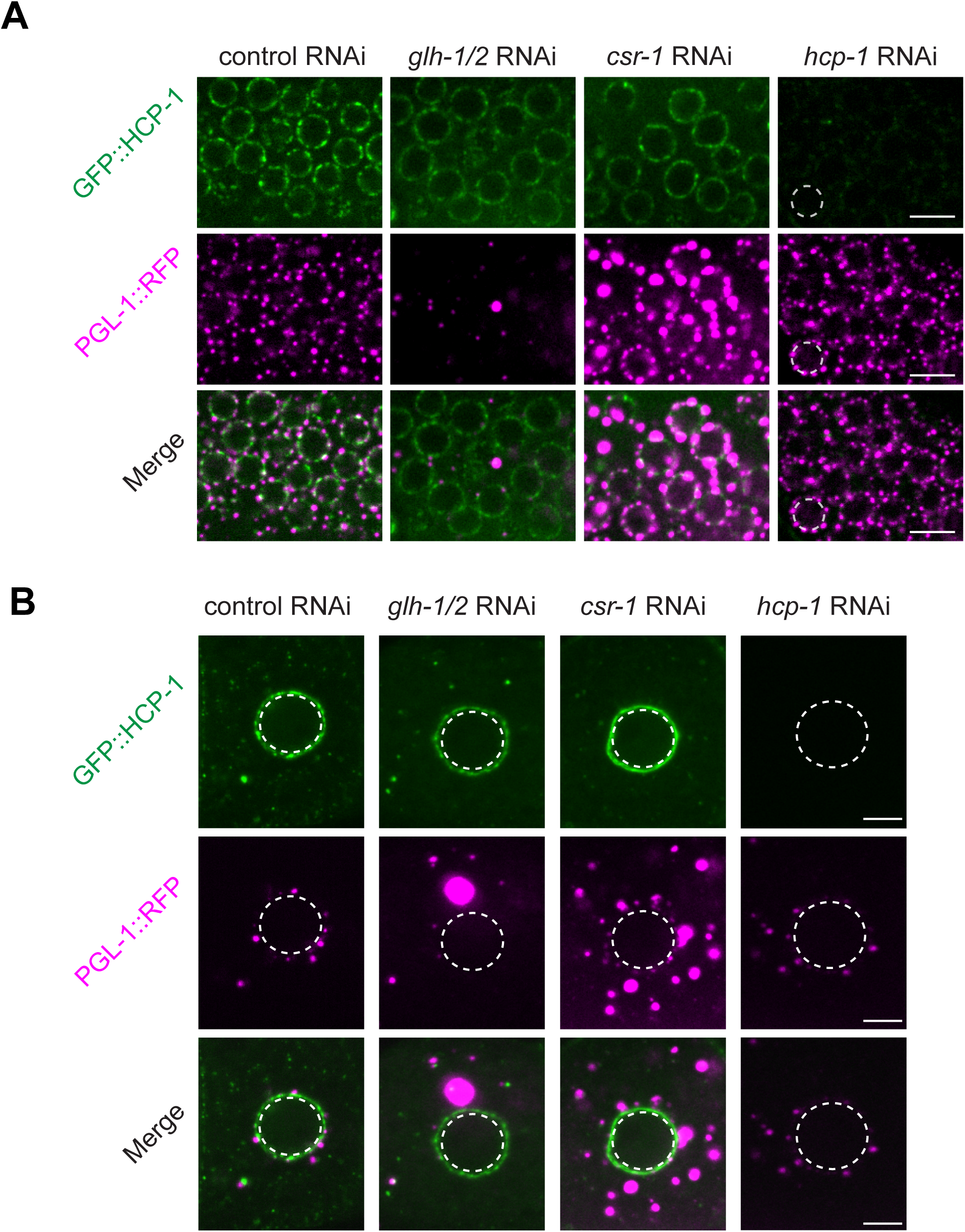
Formation of HCP-1 foci and P granules are independent. A. Single-plane images (60x objective) of GFP::HCP-1 and PGL-1::RFP at the mitotic zone treated with control (L4440), *glh-1/glh-2*, *csr-1*, or *hcp-1* RNAi, respectively. White dashed circle outlines a single nucleus. Scale bar: 5 µm. **B.** Single-plane images (60x objective) of GFP::HCP-1 and PGL-1::RFP at an oocyte treated with control (L4440), *glh-1/glh-2*, *csr-1*, or *hcp-1* RNAi, respectively. White dashed circles depict the nuclear periphery. Scale bar: 5 µm.

We next performed reciprocal experiments in which *hcp-1* was depleted in the same reporter strain. The RNAi treatment led to at least three-fold reduction in GFP::HCP-1 signals (Figure S3). Despite this loss, PGL-1::RFP still formed robust perinuclear granules similar to wild-type (Figures 3A and 3B). Together, these results indicate that HCP-1 foci and P granules form independently and likely represent distinct cytoplasmic compartments.

### NPP-14 is required for HCP-1 foci formation

Because HCP-1 colocalizes with nuclear pore proteins (Figure 1), we asked whether the formation of HCP-1 foci and nucleoporin foci are interdependent. To test whether nucleoporin foci require HCP-1, we examined NPP-11::wrmScarlet in *hcp-1* deletion mutants where the *hcp-1* open reading frame was deleted by CRISPR genome editing. In the absence of HCP-1, NPP-11 still formed clusters at germline nuclei and cytoplasmic foci in oocytes, similar to wild-type animals (Figures 4A and 4B), indicating that HCP-1 is not required for the assembly of nucleoporin foci.

**Figure 4.**
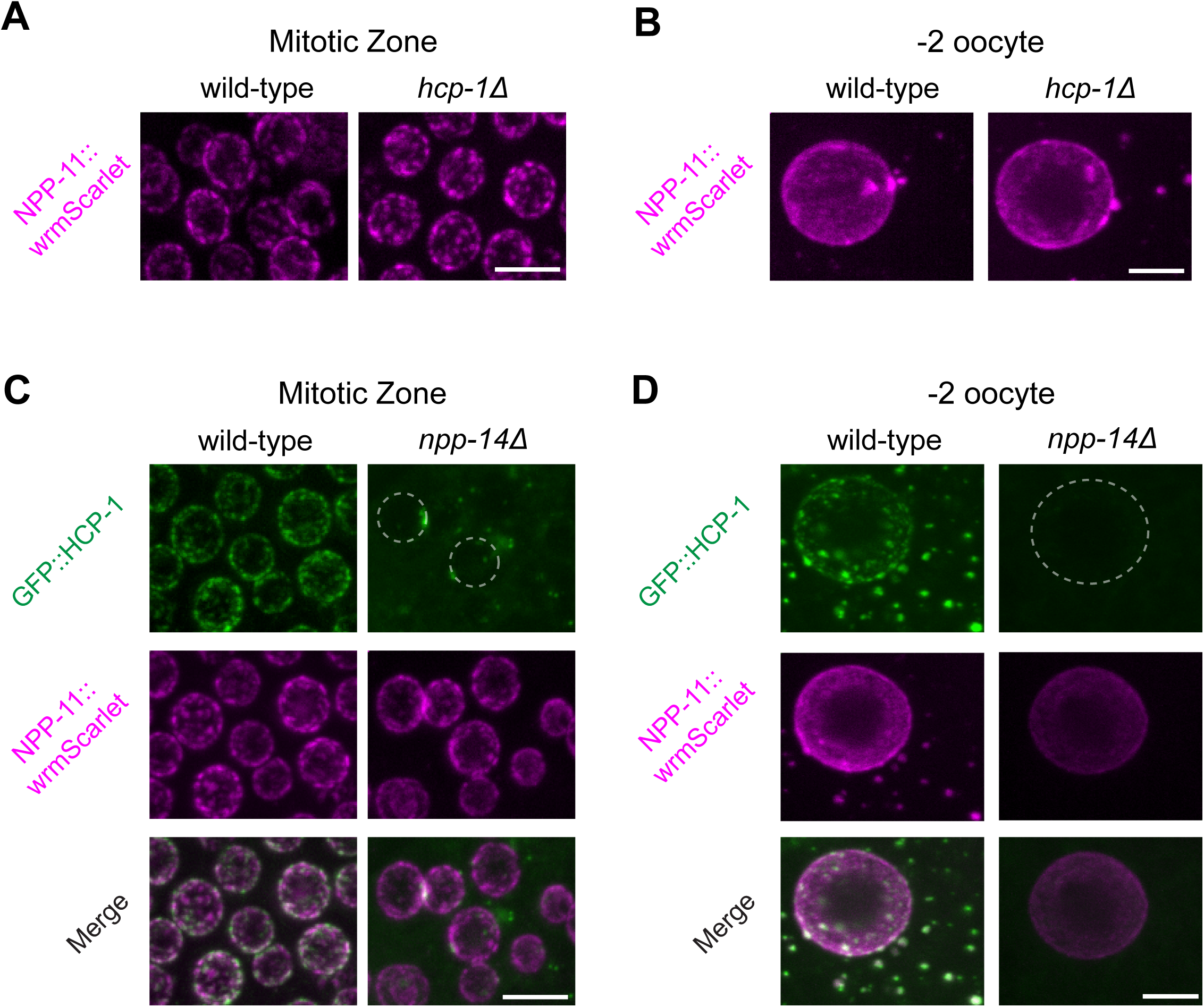
NPP-14 is required for HCP-1 foci formation. **A.** Maximum intensity projections (60x objective) of germ nuclei at the mitotic zone expressing NPP-11::wrmScarlet from wild-type or *hcp-1(how54)* worms. Scale bar: 5 µm. **B.** Maximum intensity projections (60x objective) of -2 oocytes expressing NPP-11::wrmScarlet from wild-type or *hcp-1(how54)* worms. Scale bar: 5 µm. **C.** Maximum intensity projections (60x objective) of germ nuclei at the mitotic zone expressing GFP::HCP-1 and NPP-11::wrmScarlet from wild-type or *npp-14(how55)* worms. White dashed circles outline nuclei. Scale bar: 5 µm. **D.** Maximum intensity projections (60x objective) of -2 oocytes expressing GFP::HCP-1 and NPP-11::wrmScarlet from wild-type or *npp-14(how55)* worms. White dashed circle outlines the nucleus. Scale bar: 5 µm.

We next performed the reciprocal experiment to test whether nucleoporins are required for HCP-1 foci formation. While most nucleoporins are essential for viability, NPP-14 (a component of the nuclear pore cytoplasmic filaments) is dispensable for nuclear pore assembly and *C. elegans* survival (Thomas et al., 2023). We therefore introduced a null allele of *npp-14* into a strain co-expressing GFP::HCP-1 and NPP-11::wrmScarlet. Consistent with previous findings (Thomas et al., 2023; Thomas et al., 2025), loss of NPP-14 abolished nucleoporin foci in oocytes, although NPP-11 remained localized at the nuclear envelope (Figures 4C and 4D). Deletion of *npp-14* did not significantly reduce overall GFP::HCP-1 protein levels relative to wild type (Figure S4), but it led to a depletion of perinuclear GFP::HCP-1 and a complete loss of cytoplasmic HCP-1 foci in oocytes (Figures 4C and 4D). Together, these results suggest that while HCP-1 is dispensable for nucleoporin foci assembly, NPP-14 is required for the formation of HCP-1 foci.

### HCP-1 translocates from the nuclear envelope to the nucleus during mitosis

Next, we sought to determine the physiological function of perinuclear HCP-1 in germ cell nuclei. We hypothesized that HCP-1 is stored at the nuclear periphery and imported into the nucleus during mitosis. In the *C. elegans* germline, only a small number of germ cells undergo mitosis at any given time (Maciejowski et al., 2006). Careful examination of the mitotic region revealed that on average 2.5 nuclei contain nuclear GFP::HCP-1 (Figures 5A and 5B), indicating import of HCP-1 to the nucleus.

**Figure 5.**
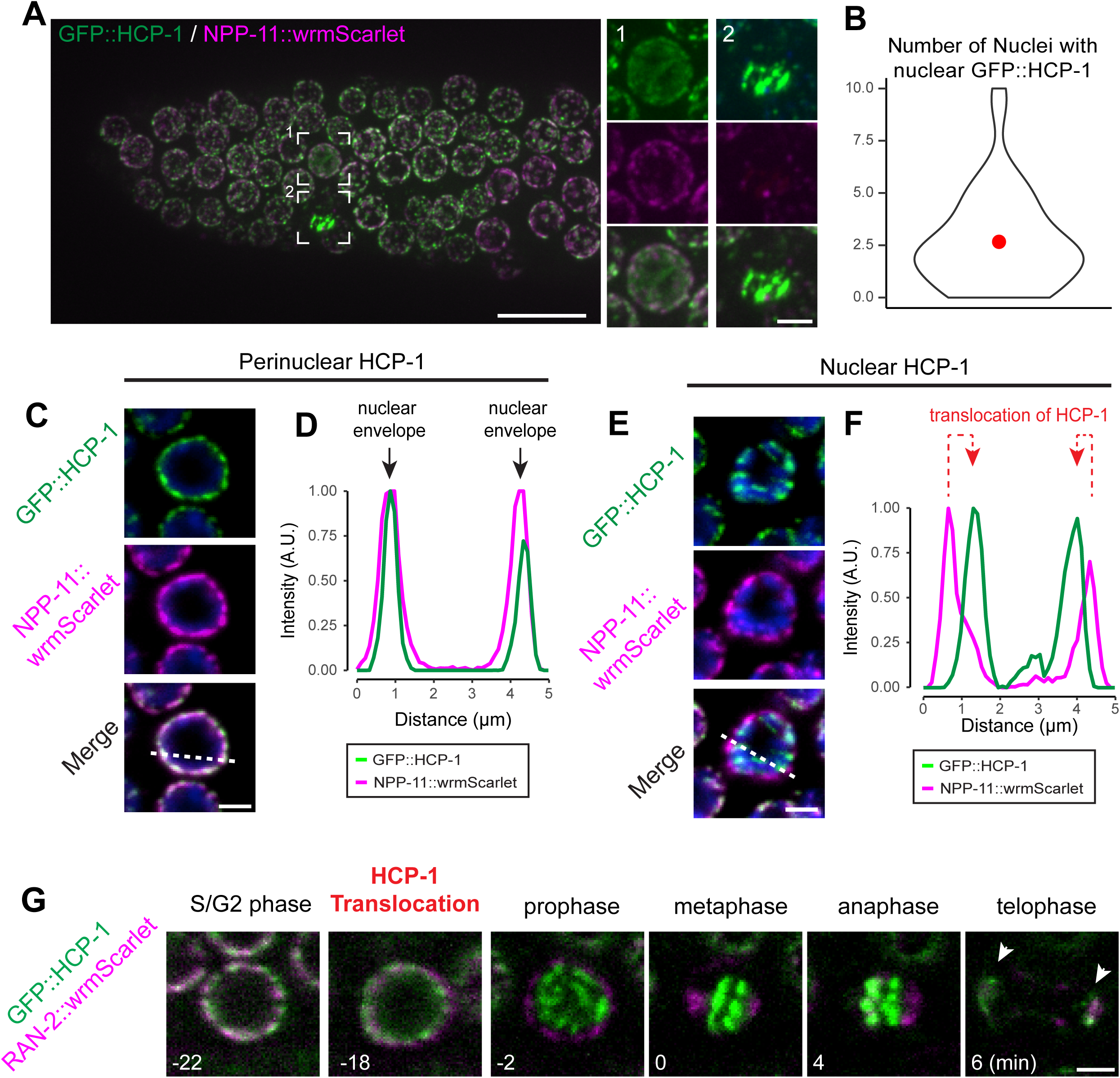
Perinuclear HCP-1 translocates into nuclei during mitosis. **A.** Maximum intensity projection (60x objective) of the mitotic zone of a dissected gonad expressing GFP::HCP-1 and NPP-11::wrmScarlet (scale bar: 10 µm). Dashed squares indicate insets to the right (scale bar: 2 µm) showing HCP-1 (1) diffuse nuclear signal or (2) condensed nuclear signal. **B.** Violin plot showing the number of nuclei with nuclear GFP::HCP-1 at the mitotic zone for each day 2 adult gonad (n=42). Red dot indicates the mean. **C.** Representative fixed, single-plane image of a nucleus at the mitotic zone expressing GFP::HCP-1 and NPP-11::wrmScarlet, showing perinuclear HCP-1 localization. White dashed line indicates the linescan in D. Scale bar: 2 µm. **D.** Intensity profile of GFP::HCP-1 (green) or NPP-11::wrmScarlet (magenta) along the dashed line in C. Arrows indicate signal at the nuclear envelope. **E.** Representative fixed, single-plane image of a nucleus at the mitotic zone expressing GFP::HCP-1 (green) and NPP-11::wrmScarlet (magenta), showing nuclear HCP-1 localization at condensed chromosomes. White dashed line indicates the linescan in F. Scale bar: 2 µm. **F.** Intensity profile of GFP::HCP-1 (green) or NPP-11::wrmScarlet (magenta) along the dashed line in E. Red dashed arrows indicate the translocation of HCP-1 showing nuclear HCP-1 signal (green) within the nuclear envelope (magenta). **G.** Single-plane timelapse (60x objective) of a nucleus at the mitotic zone expressing GFP::HCP-1 and RAN-2::wrmScarlet throughout mitosis in a L4 worm. Timing in minutes (min) is shown relative to metaphase (0). White arrows indicate the daughter nuclei formed in telophase. Scale bar: 2 µm.

We employed two independent approaches to assess whether HCP-1 translocates from the nuclear envelope into the nucleus during cell division. In the first approach, we fixed animals co-expressing GFP::HCP-1 and NPP-11::wrmScarlet and stained with DAPI to visualize DNA. As expected, most germ cell nuclei displayed colocalization of HCP-1 and NPP-11 at the nuclear envelope (Figures 5C and 5D). However, we also identified nuclei in which perinuclear HCP-1 signal was lost and GFP::HCP-1 instead accumulated on condensed chromosomes (Figures 5E and 5F). Notably, in these nuclei the nuclear envelope, as marked by NPP-11::wrmScarlet, remained intact (Figure 5E), suggesting that HCP-1 translocation precedes complete nuclear envelope breakdown. In the second approach, we performed live time-lapse confocal microscopy in animals expressing GFP::HCP-1 and RAN-2::wrmScarlet, a Ran GTPase-activating protein that localizes to the nuclear periphery and regulates nucleocytoplasmic transport (Thomas et al., 2023). Germline nuclei were tracked over a 40-min imaging window. Cells undergoing mitosis were identified and further analyzed. Because HCP-1 associates with condensed chromosomes at metaphase, we defined the time of chromosome alignment at the metaphase plate as time point 0. Between -22 and -18 minutes, we observed the redistribution of GFP::HCP-1 from the nuclear envelope into the nuclear interior with continued presence of RAN-2::wrmScarlet at the nuclear periphery (Figure 5G).

To determine whether HCP-1 is actively imported into the nucleus or passively translocated during partial nuclear envelope breakdown, we searched for potential nuclear localization signals (NLSs) using cNLS Mapper prediction software (Kosugi et al., 2009). This analysis identified a putative bipartite NLS at the N-terminus of HCP-1 (Figure S5A). To test its function, we used CRISPR/Cas9 genome editing to delete the sequences encoding predicted NLS from the endogenous *hcp-1* locus. However, mutated GFP::HCP-1 still translocated into the nucleus in both germ cells and embryos (Figures S5B and S5C), indicating that this sequence is dispensable for nuclear entry. Although a canonical NLS could not be identified, our findings suggest HCP-1 is translocated to the nucleus either prior to or during nuclear envelope breakdown in germ cells.

### NPP-14 is required for germ cell proliferation and embryonic cell division

Because NPP-14 promotes the enrichment of perinuclear HCP-1 (Figure 4) and perinuclear HCP-1 subsequently translocates to nuclei in the mitotic region (Figure 5), we investigated whether NPP-14 influences HCP-1 accumulation in dividing germ cells. Indeed, the nuclear GFP::HCP-1 signal in *npp-14* mutant animals was significantly lower than that in wild-type (Figures 6A and 6B). In addition, we found that *npp-14* mutant gonads were smaller and contained fewer germ nuclei when compared to wild type (Figures 6C and 6D), indicating that NPP-14 is essential for germ cell proliferation.

**Figure 6.**
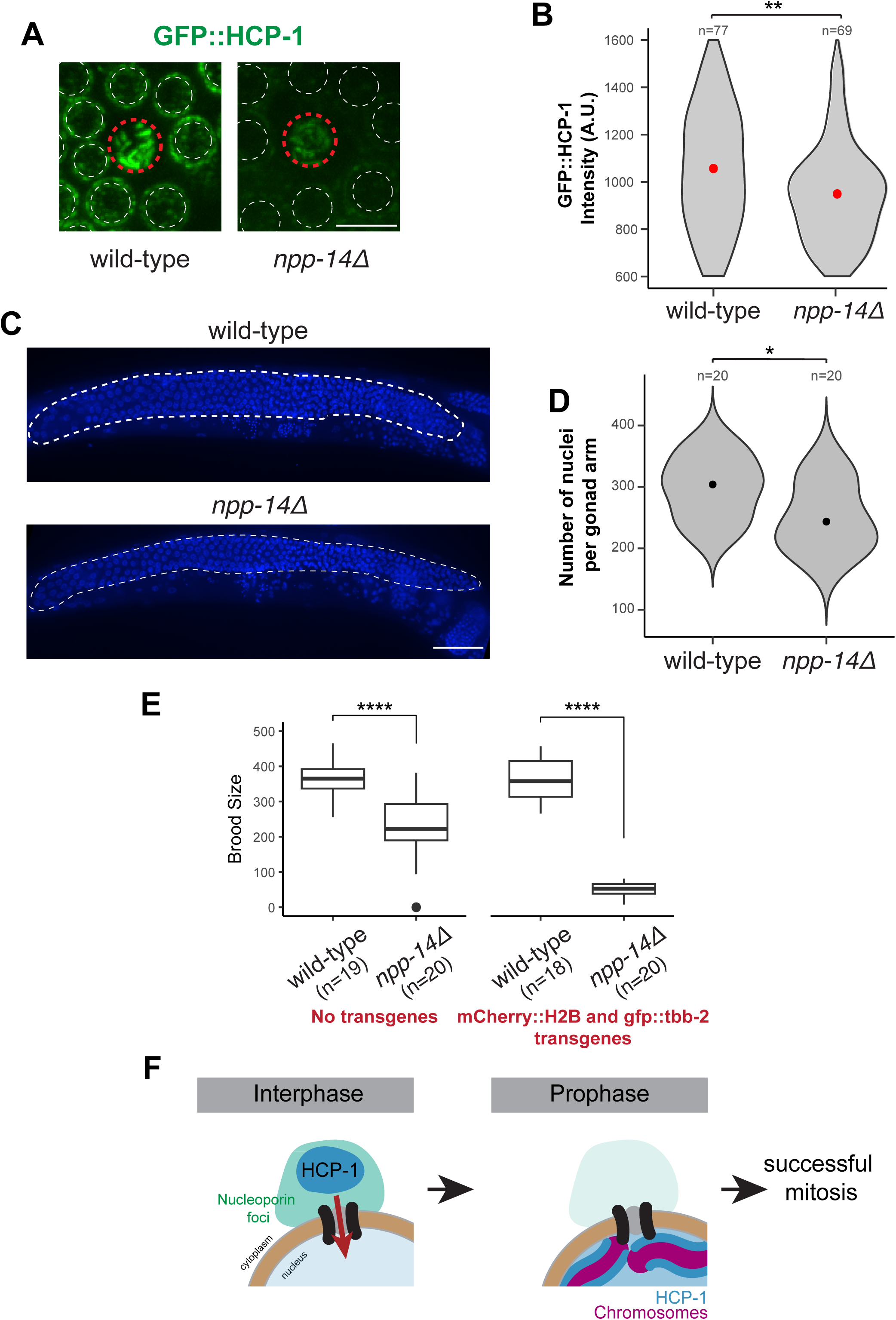
NPP-14 promotes germ cell division and fertility. **A.** Maximum intensity projections (60x objective) of germ nuclei at the mitotic zone expressing GFP::HCP-1 from wild-type or *npp-14(ax4543)* worms, showing reduced nuclear HCP-1 in *npp-14* mutants. Red dashed circles outline a nucleus in prophase with nuclear HCP-1 as quantified in B. White dashed circles depict the nuclear periphery for nuclei with perinuclear HCP-1. Scale bar: 5 µm. **B.** Violin plot showing nuclear GFP::HCP-1 intensity of M-phase germ nuclei at the mitotic zone from wildtype or *npp-14(ax4543)* worms. Red dot indicates the mean. *n* refers to the number of nuclei analyzed. Statistics determined using Wilcoxon rank-sum test and ** represents *p* < 0.01. **C.** Maximum intensity projections (40x objective) of a wild-type or *npp-14(ax4543)* adult carrying the *mCherry::h2b*; *gfp::tbb-2* transgene, with the germline visualized by DAPI staining. Scale bar: 40 µm. **D.** Violin plot showing number of nuclei per gonad arm surface of wild-type or *npp-14(ax4543)* adults carrying the *mCherry::h2b*; *gfp::tbb-2* transgene. Black dots represent the median. *n* refers to the number of worms analyzed. Statistics determined using Wilcoxon rank-sum test and * represents p<0.05. **E.** Boxplot showing median fertility of wild-type or *npp-14(ax4543)* worms in the N2 background (left) or carrying the *mCherry::h2b*; *gfp::tbb-2* transgene (right) at 20°C. *n* refers to the number of worms assayed. Statistics determined using Wilcoxon rank-sum test and **** represents p<0.0001. **F.** Model showing the dual localization of HCP-1, which localizes to nucleoporin foci during interphase and translocates to the nucleus during prophase to promote mitosis in the *C. elegans* germline.

To assess whether these germline defects affect fertility, we measured brood size. Under normal growth conditions, wild type strains produced ∼363 progeny per animal, while *npp-14* mutants produced ∼227 progeny (Figure 6E). Notably, in a genetic background expressing translational fusions of GFP to β-tubulin (GFP::TBB-2) and mCherry to Histone (mCherry::H2B) (Connolly et al., 2015), the brood size was reduced from ∼357 progeny in wild type to ∼51 in *npp-14* mutant strains (Figure 6E).

To determine the underlying cause of this reduced fertility and the effect of this presumably sensitized background, we examined embryonic lethality phenotype. Without transgene, both wild-type and *npp-14* mutant animals showed close to 100% viable embryos (Figure S6A) (Thomas et al., 2023). However, the expression of GFP::TBB-2 and mCherry::H2B resulted in ∼21% embryo lethality in *npp-14* mutants (Figure S6A). Since GFP::TBB-2 and mCherry::H2B mark microtubules and chromosome respectively (Connolly et al., 2015), we performed spinning-disk confocal microscopy to monitor spindle assembly and chromosome segregation. Because fluorescent signals were too weak to quantify in the mitotic germline region, we compared spindle assembly in live embryos. Wild-type embryos, as expected, assembled only bipolar spindles during mitotic division (Figures S6B and S6C). However, a significant fraction (∼40%) of *npp-14* mutant embryos assembled multipolar spindles in the first cell cycles (Figures S6B and S6C). Live imaging further confirmed the defects in spindle assembly and chromosome segregation in *npp-14* mutants (Figure S6D). Together, our findings suggest that NPP-14 is required for normal germ cell proliferation and nuclear HCP-1 accumulation, and its loss leads to significant mitotic defects in the early embryo in a sensitized genetic background.

## Discussion

In this study, we uncover a previously unrecognized spatial regulation of the kinetochore component HCP-1 in the *C. elegans* germline (Figure 6F). While kinetochore proteins are generally thought of as nuclear factors required for chromosome segregation, our findings demonstrate that HCP-1 forms distinct perinuclear and cytoplasmic condensates in germ cells, which are spatially and functionally distinct from P granules. HCP-1 granules are particularly enriched in mitotic nuclei and oocytes, suggesting a developmental regulation of its localization and function.

We propose a model in which nuclear pore proteins organize perinuclear HCP-1 compartments that act as reservoirs for kinetochore components. Upon mitotic entry, HCP-1 is released from these condensates and translocated into the nucleus to support kinetochore assembly (Figure 6F). Specifically, we show that perinuclear HCP-1 colocalizes with nuclear pore proteins, such as NPP-11 and NPP-14. The strong colocalization between HCP-1 and these NPPs suggests that nuclear pore complexes may serve as organizational hubs for kinetochore component sequestration or regulation (Figure 6F). Importantly, we demonstrate that NPP-14 is required for the formation of HCP-1 condensates, while HCP-1 is dispensable for the assembly of nucleoporin foci, suggesting a hierarchy assembly. Our data further reveal that perinuclear HCP-1 is dynamically translocated into the nucleus during mitosis, where it contributes to kinetochore assembly and chromosome segregation. This translocation appears to occur prior to complete nuclear envelope breakdown, suggesting a regulated import mechanism rather than passive diffusion. Future studies should identify the molecular scaffold that directly recruits HCP-1 and clarify how HCP-1 transitions from perinuclear storage to nuclear import.

Functionally, the loss of NPP-14 impairs the formation of HCP-1 granules, reduces nuclear HCP-1 accumulation during mitosis, and leads to defects in germ cell proliferation and embryonic cell division. These phenotypes are accompanied by reduced brood size and increased embryonic lethality, particularly in sensitized genetic backgrounds. Notably, a significant proportion of *npp-14* mutant embryos exhibit multipolar spindle formation, implicating NPP-14 in maintaining spindle integrity and proper chromosome segregation, through mechanisms dependent or independent of HCP-1.

The perinuclear localization of HCP-1 in *C. elegans* germ cells echoes observations in vertebrate systems, where the human homolog CENP-F is localized to the nuclear envelope during interphase and recruited by Nup133 (Berto et al., 2018). However, the formation of cytoplasmic condensates and the dependency on nuclear pore components such as NPP-14 appear to be unique features of the *C. elegans* germline. These findings raise intriguing questions about the evolutionary conservation and divergence of kinetochore regulation mechanisms across species.

In summary, our study reveals a novel role for nuclear pore proteins in organizing kinetochore components in the germline (Figure 6F). The discovery of perinuclear and cytoplasmic HCP-1 condensates expands our understanding of kinetochore biology and highlights the importance of spatial compartmentalization in regulating cell division. Future studies will be needed to elucidate the molecular mechanisms underlying HCP-1 condensation and its functional consequences for germline development and chromosome segregation.

## Materials and Methods

### Maintenance of *C. elegans* strains

The *C. elegans* wild-type strain is N2 unless otherwise indicated, and all strains are in the N2 Bristol background (Brenner, 1974). Worms were cultured according to standard growth conditions at 20°C on Nematode Growth Media (NGM) seeded with OP50-1 *E. coli*. All strains reported in this study are listed in Table S1.

### CRISPR/Cas9 genome editing

CRISPR/Cas9 genome editing was conducted by microinjection as previously reported (Ghanta and Mello, 2020; Paix et al., 2015). To generate *hcp-1(how54)*, the open reading frame (ORF) was removed using two guide RNAs in equimolar amounts (34 µM total) flanking the ORF with a single-stranded donor (ssdonor, 1 µg/µL) as a repair template. The ssdonor included 35 nt homology arms upstream and downstream of the ORF and a HINDIII restriction site for genotyping. *npp-14(how55)* ORF deletion was generated using two guide RNAs flanking the ORF, but without a single-stranded donor. *hcp-1(how56)* nuclear localization signal (NLS) mutants were generated in a similar fashion, using two guide RNAs flanking the predicted NLS with a ssdonor with homology arms flanking the predicted NLS as a repair template. Guide RNAs and donor sequences are listed in Table S2. The PRF4 vector containing *rol-6* was used as a co-injection marker (Ghanta and Mello, 2020).

### Microscopy

All images of adult germlines were conducted approximately 48 hours post L4 (day 2 adults). Timelapse images in Figure 5 were conducted at the L4 stage as M-phase germline cells are more frequently observable at this stage.

For live imaging, day 2 adults were paralyzed with 5 mM levamisole and mounted on glass slides with 4% agar pads. For imaging of dissected gonads, day 2 adults were dissected into M9 on a ring slide and immediately imaged. To image live embryos, adults were dissected into M9 and embryos transferred to glass slides with 4% agar pads. To image fixed adults, worms were fixed via methanol/acetone fixation. In brief, worms were fixed in cold methanol, then cold acetone, 20 mins each at -20°C. The animals were gradually rehydrated with 75%, 50%, then 25% acetone washes, 15 mins each, at -20°C. Fixed worms were resuspended in Vectashield antifade mounting medium with DAPI (Vector laboratories) for imaging.

All imaging was conducted with a Nikon Ti2 inverted microscope equipped with an X-Light V3 spinning disk confocal unit (CrestOptics) in NIS-Elements AR 5.41.02, apart from FRAP images. The Plan Apo 40x objective was used to image whole germlines; Plan Apo 60x oil objective to image enlarged views of the germline, oocytes, embryos, and timelapse images. Imaging for FRAP was conducted using a Zeiss Axio Observer microscope with an Airyscan two detector using the Plan Apo 63x objective. FRAP images were processed using standard Airyscan processing. All images were further processed using FIJI.

### Image Quantification

To quantify the degree of colocalization between HCP-1 and either PGL-1 or NPP-11, a region of interest (ROI) centered and encompassing a single nucleus of a single-plane image was defined for at least fourteen randomly selected nuclei in the mitotic region of the germline for each strain. Pearson’s correlation coefficient for the two channels was calculated for each ROI using the Jacop2 plugin in FIJI.

To quantify the roundness of HCP-1 foci, ROIs were drawn in the mitotic region of the germline for each of seven randomly selected worms or spanning at least 3 oocytes for each of eleven randomly selected worms, using single-plane images. Pixel intensities between 30-80% of the maximum were included to avoid saturated or background signals. Foci with areas between 0.2-1 µm^2^ were selected, and roundness was calculated from the 2D thresholded images using the Shape Descriptor tool in ImageJ. Roundness was calculated as follows, with 1 representing a perfect circle and 0 representing an elongated irregular object: Roundness 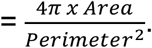

To quantify nuclear GFP::HCP-1 intensity, M-phase nuclei (as indicated by condensed chromosomes) at the mitotic zone were selected. Circular ROIs (14 µm^2^) were drawn on Z-stack images spanning 2.7 µm (0.3 µm step size) to encompass the entire nuclear HCP-1 signal. Mean nuclear GFP intensity was measured in FIJI, subtracting background GFP fluorescence outside the worm. At least 33 *gfp::hcp-1* or *gfp::hcp-1*; *npp-14(ax4543)* day 2 adults were analyzed.

### Fluorescence Recovery after Photobleaching (FRAP)

Photobleaching of HCP-1 foci in the mitotic region of the germline or oocytes in live day 2 adults was performed using 100% laser power and 25 iterations in the 488 nm channel with circular ROIs slightly larger than the foci. A single-plane image was taken at 4 timepoints prior to photobleaching, and subsequently every 10 seconds after photobleaching.

Images were processed via standard Airyscan processing (AS 3.9) and further analyzed in FIJI. For each timepoint, the fluorescence intensity (I_t)_ of the bleached foci was corrected for background intensity outside of the worm (I_back_), then normalized to the average initial fluorescence intensity (I_0_) of the bleached foci corrected to initial background intensity outside of the worm (I_0,back_) prior to photobleaching of 4 timepoints to calculate relative HCP-1 intensity (I_norm_): 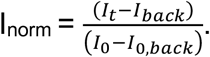

To calculate the recovery of HCP-1 after photobleaching, a first order exponential model was fit to individual FRAP curves, offset to the mean intensity of the bleached foci at photobleaching at time zero (y_0_), where A_rec_ is the fluorescence recovery amplitude and *k* is the recovery rate constant: I= y_0_ + A_rec_ * (1-e^-***k***Time^). t_1/2_ was calculated from the first order exponential model. The mobile fraction was defined as the percentage of HCP-1 that can recover from the baseline bleached intensity (y_0_) and calculated as follows: Mobile Fraction 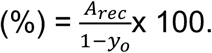

### RNAi by feeding

RNAi was conducted by feeding *C. elegans* with HT115(DE3) *E. coli* expressing the designated RNAi vector. Empty RNAi vector L4440, *csr-1*, *hcp-1* RNAi bacteria strains were sourced from the Ahringer RNAi Collection (Source Biosciences); and *glh-1/2* RNAi strain from Updike et al.,(Updike et al., 2014). NGM agar plates supplemented with 50 µg/ml ampicillin and 5 mM IPTG were seeded with RNAi bacterial cultures and incubated at room temperature for at least 2 days to induce dsRNA production prior to feeding. For RNAi feeding, L4s expressing GFP::HCP-1; PGL-1::RFP were plated on the supplemented NGM and F1 day 2 adults (treated one generation) were imaged to assay HCP-1 and P granule localization. Because animals treated one generation with *csr-1* RNAi displayed severe fertility defects, RNAi treatment for *csr-1* began with the L1 stage and day 2 adult worms (from the same generation) were imaged.

### Quantification of number of germ cells

Synchronized wild-type or *npp-14(ax4543)* adults carrying the *mcherry::h2b* and *gfp::tbb-2* transgenes were fixed by cold methanol/acetone and germlines visualized by DAPI staining. To quantify number of germ nuclei per gonad arm, 20 animals were selected for which the whole germline was visible, and the images converted to Z-projections spanning the top layer of germline nuclei (4.8 µm, step=1.2 µm). Germline nuclei were manually counted in each germline arm using the multipoint tool in FIJI.

### Brood Size analyses

For wild-type, *npp-14(ax4543)* and wild-type, *npp-14(ax4543)* in the *mcherry::*h2b and *gfp::tbb-*2; transgene background, 18 L4s at minimum were single picked to NGM plates and grown at 20°C. After two days, the mothers were transferred to fresh NGM plates, and progeny counted at the end of egg-laying. The brood size for each animal was calculated by adding the progeny from the original and transferred plates.

### Network analyses

Proteins enriched in *glh-1::turboID* were analyzed using STRING (string-db.org) with high confidence (Szklarczyk et al., 2023), then exported to Cytoscape to examine functional enrichment of network clusters (Shannon et al., 2003). Singletons were excluded from the network.

### Western blot analyses

To quantify HCP-1 abundance, 1600 synchronized *GFP::hcp-1* or *GFP::hcp-1*; *npp-14(ax4543*) L1s were grown to adulthood, with two independent biological replicates for each strain. Adults were washed twice with M9, resuspended in protein loading buffer (85 μL Tris-HCl pH 8.0, 5 μL 1 M DTT, 30 μL 4x LDS loading dye), and boiled for 10 min. Samples were run on a precast SDS-PAGE gel (Novex, Invitrogen) and transferred to nitrocellulose membrane (GE Healthcare Life Sciences). After blocking with 3% milk in PBST, the membranes were incubated at 4°C overnight with either mouse-*anti*-GFP (Roche, 1:800) or rat-*anti*-tubulin (Bio-rad, 1:1000). Membranes were then washed with PBST, subsequently incubated with either goat-*anti*-mouse IgG H&L HRP (Abcam) or goat-*anti*-rat IgG H&L HRP (Abcam). Membranes visualized with Max ECL substrate (HCP-1) or ECL substrate (tubulin).

### Embryonic Lethality

For wild-type, *npp-14(ax4543)*, and their counterparts carrying the *mcherry::*h2b; gfp*::tbb-2* transgene, ∼50 embryos in triplicates were transferred to NGM. After two days, the numbers of hatched and unhatched (dead) embryos were counted. Dead embryos (%) was calculated as follows: Dead embryos 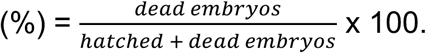

## Acknowledgement

We thank members in the Tang lab for discussion and critical comments, Dr. J. Dumont for sharing GFP::HCP-1 strains, Dr. G. Seydoux for sharing NPP related strains. J. Wagner and H. Dey for technical support, OSU Neuroscience Imaging Core for instruments (S10OD026842), the Caenorhabditis Genetics Center for providing the *C. elegans* strains (P40OD010440). I. Price was supported by Center for RNA Biology graduate fellowship and presidential fellowship at The Ohio State University. This work was supported by the Arthur Burghes professorship, NIH Maximizing Investigators’ Research Award (R35 GM142580), and NSF (MCB 2420329) to W. Tang.

---

**Figure S1.**
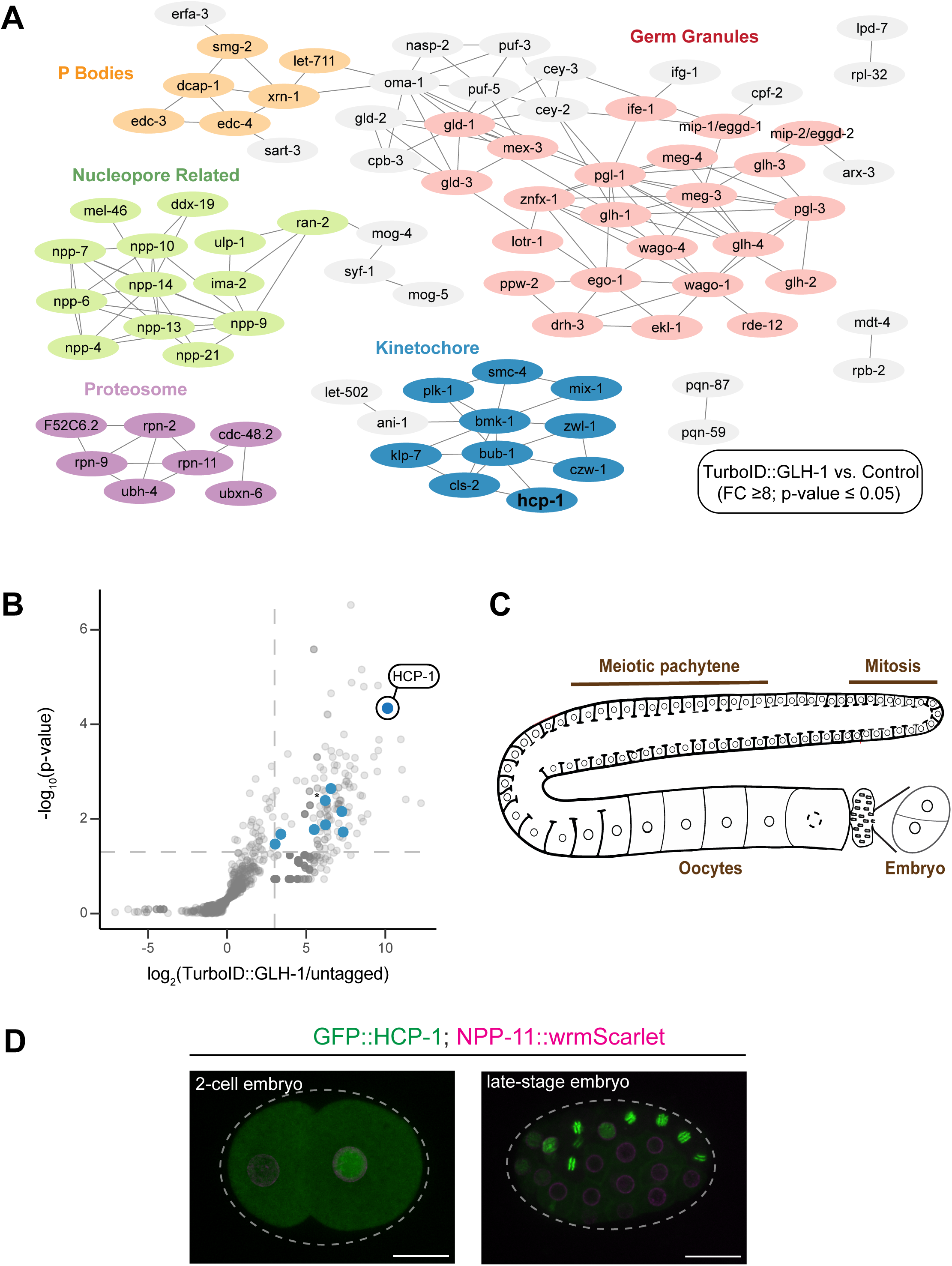
P granule proximity labeling enriches kinetochore component HCP-1. **A.** String network of proteins enriched ≥ 8-fold with TurboID::GLH-1 with a p-value ≤ 0.05 visualized in Cytoscape. Germ granule proteins shown in red, kinetochore components in blue, nucleopore related proteins in green, P-body proteins in orange, and proteosome proteins in purple. Other proteins shown in grey and singletons excluded. HCP-1 is indicated in bold. **B.** Volcano plot of enriched proteins in TurboID::GLH-1 vs untagged control. Statistics determined using a one-tailed Student’s t-test, with p < 0.05 represented by the horizontal dashed line. log2(fold change) ≥ 3 represented by the vertical dashed line. Blue dots represent the enriched kinetochore components. Asterisk indicates 2 overlapping proteins, with the most significantly enriched kinetochore component HCP-1 labeled. **C.** Schematic of *C. elegans* germline with regions of interest labeled in brown. **D.** Maximum intensity projections of GFP::HCP-1 and NPP-11::wrmScarlet in a 2-cell or late-stage embryo. Dashed lines outline each embryo. Scale bar: 15 µm.

**Figure S2.**
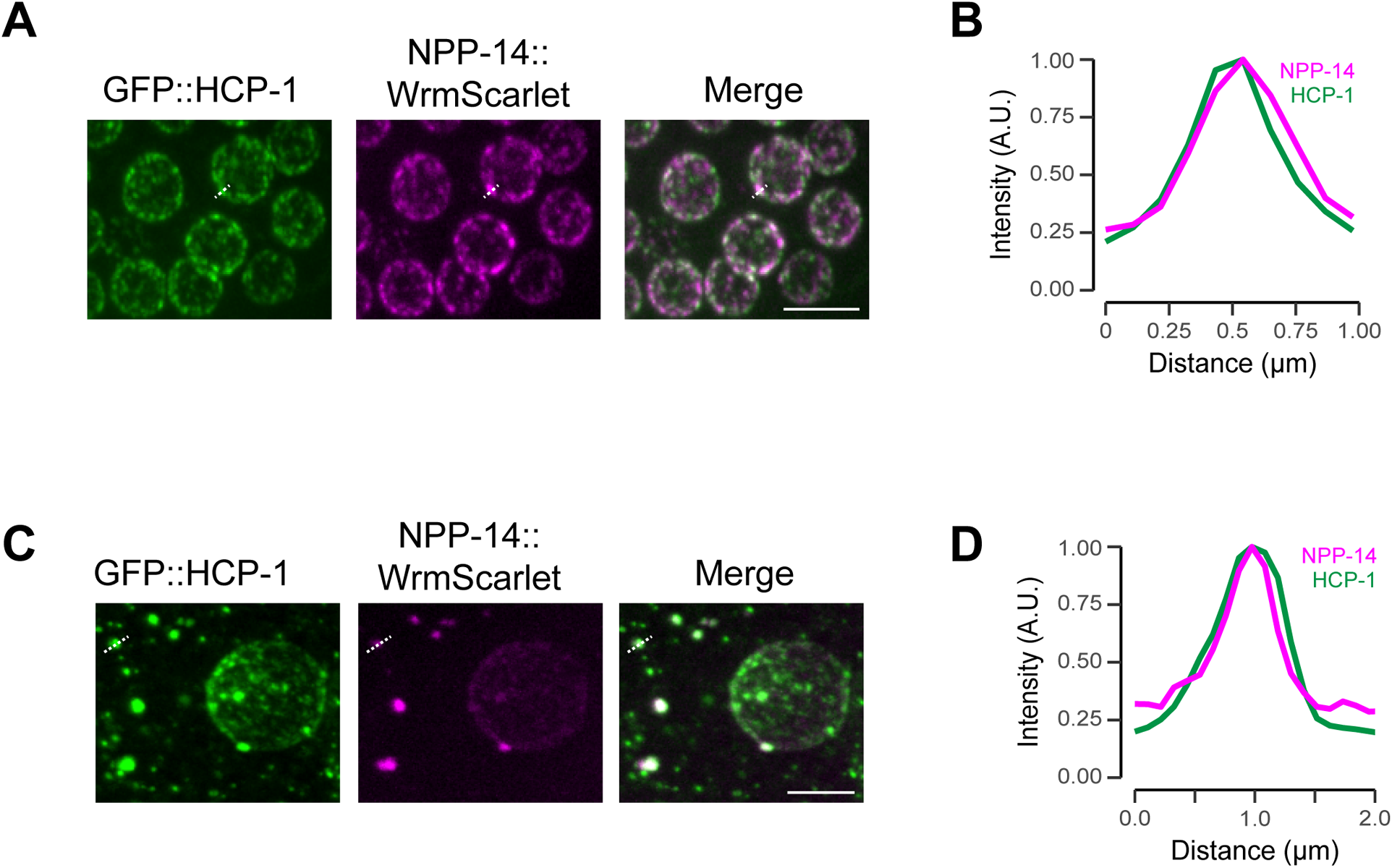
HCP-1 colocalizes with nucleoporin NPP-14. **A.** Maximum intensity projections (60x objective) of germ nuclei at the mitotic zone expressing GFP::HCP-1 and mex5p::NPP-14::wrmScarlet. White dashed line indicates the linescan in B. Scale bar: 5 µm. **B.** Intensity profile of GFP::HCP-1 (green) or NPP-14::wrmScarlet (magenta) perinuclear focus along the dashed line in A. **C.** Maximum intensity projections (60x objective) of a -2 oocyte expressing GFP::HCP-1 and mex5p::NPP-14::wrmScarlet. White dashed line indicates the linescan in D. Scale bar: 5 µm. **D.** Intensity profile of GFP::HCP-1 (green) or NPP-14::wrmScarlet (magenta) cytoplasmic focus along the dashed line in panel C.

**Figure S3.**
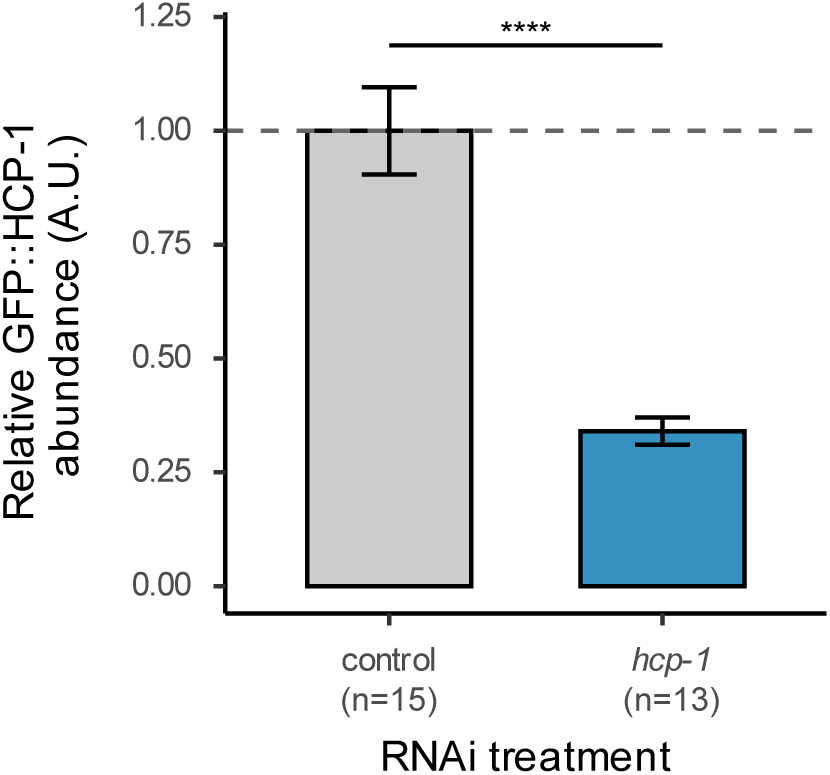
Depletion of GFP::HCP-1 using RNAi. A. Bar plot showing the average fold change of GFP::HCP-1 abundance with *hcp-1* RNAi treatment (n=13) compared to the control (n=15). *n* refers to the number of worms assayed; error bars represent standard error of the mean. Statistics determined using Wilcoxon rank-sum test and **** represents p<0.0001.

**Figure S4.**
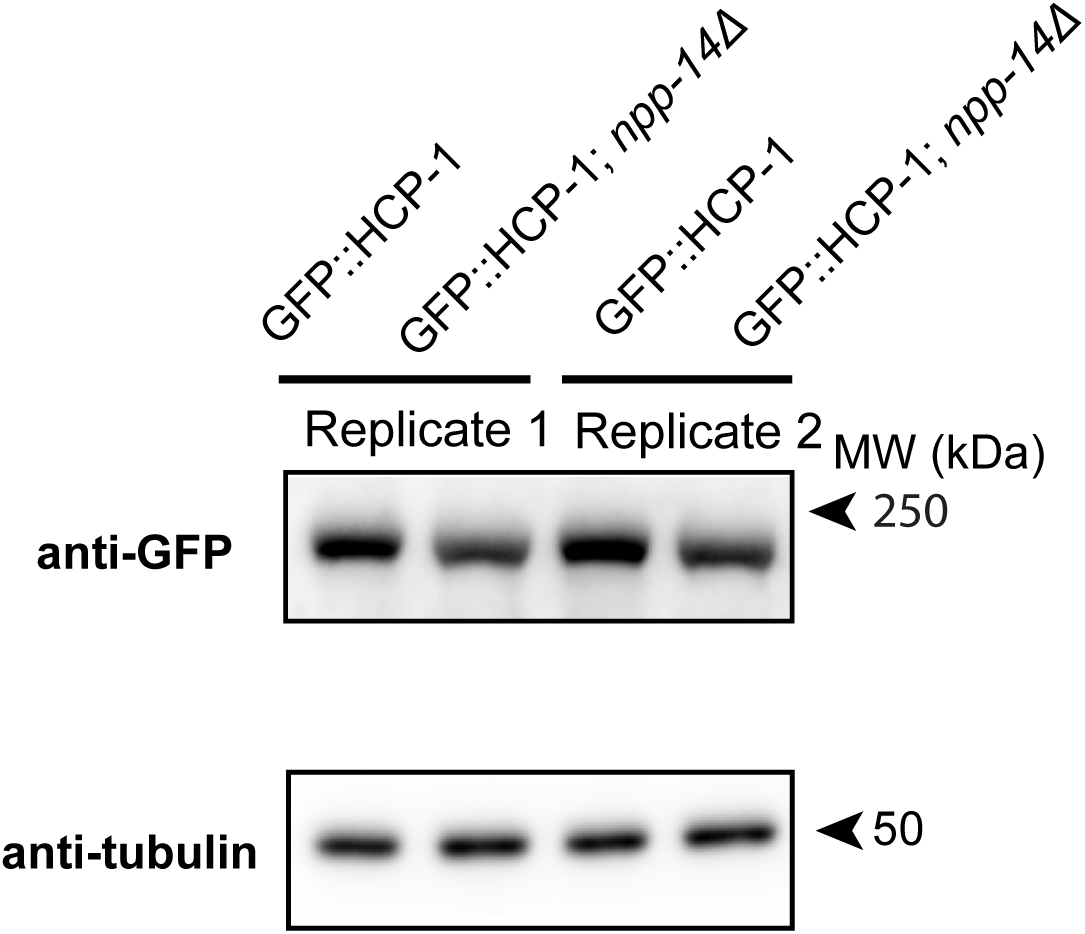
Overall HCP-1 abundance is unaffected in *npp-14* mutants. **A.** Western Blotting analysis of HCP-1 abundance in *gfp::hcp-1* and *gfp::hcp-1*; *npp-14(ax4543)* adults. HCP-1 abundance was assayed using an anti-GFP antibody with β-tubulin as the loading control.

**Figure S5.**
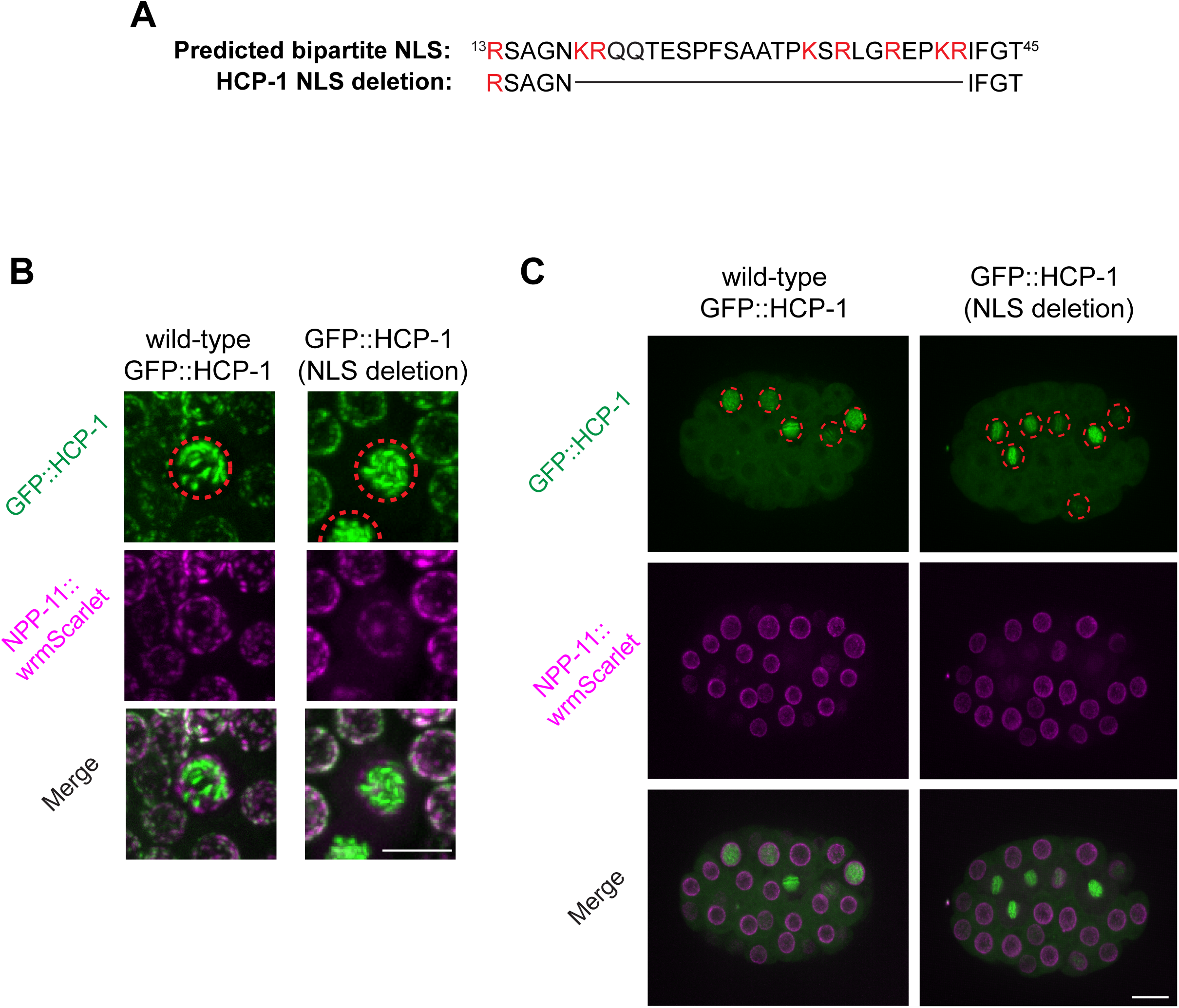
HCP-1 NLS mutants still localize to the nucleus. **A.** Schematic of predicted bipartite nuclear localization signal (NLS) and NLS deletion in *hcp-1(how56)* (bottom). Positively charged residues indicated in red. **B.** Maximum intensity projections of germ nuclei at the mitotic zone expressing GFP::HCP-1 and NPP-11::wrmScarlet in wild-type or *hcp-1(how56)* adults. Red dashed circles indicate nuclear HCP-1 localization. Scale bar: 5 µm. **C.** Maximum intensity projections of GFP::HCP-1 and NPP-11::wrmScarlet expressing wild-type or *hcp-1(how56)* embryos. Red dashed circles indicate nuclear HCP-1 signal. Scale bar: 10 µm.

**Figure S6:**
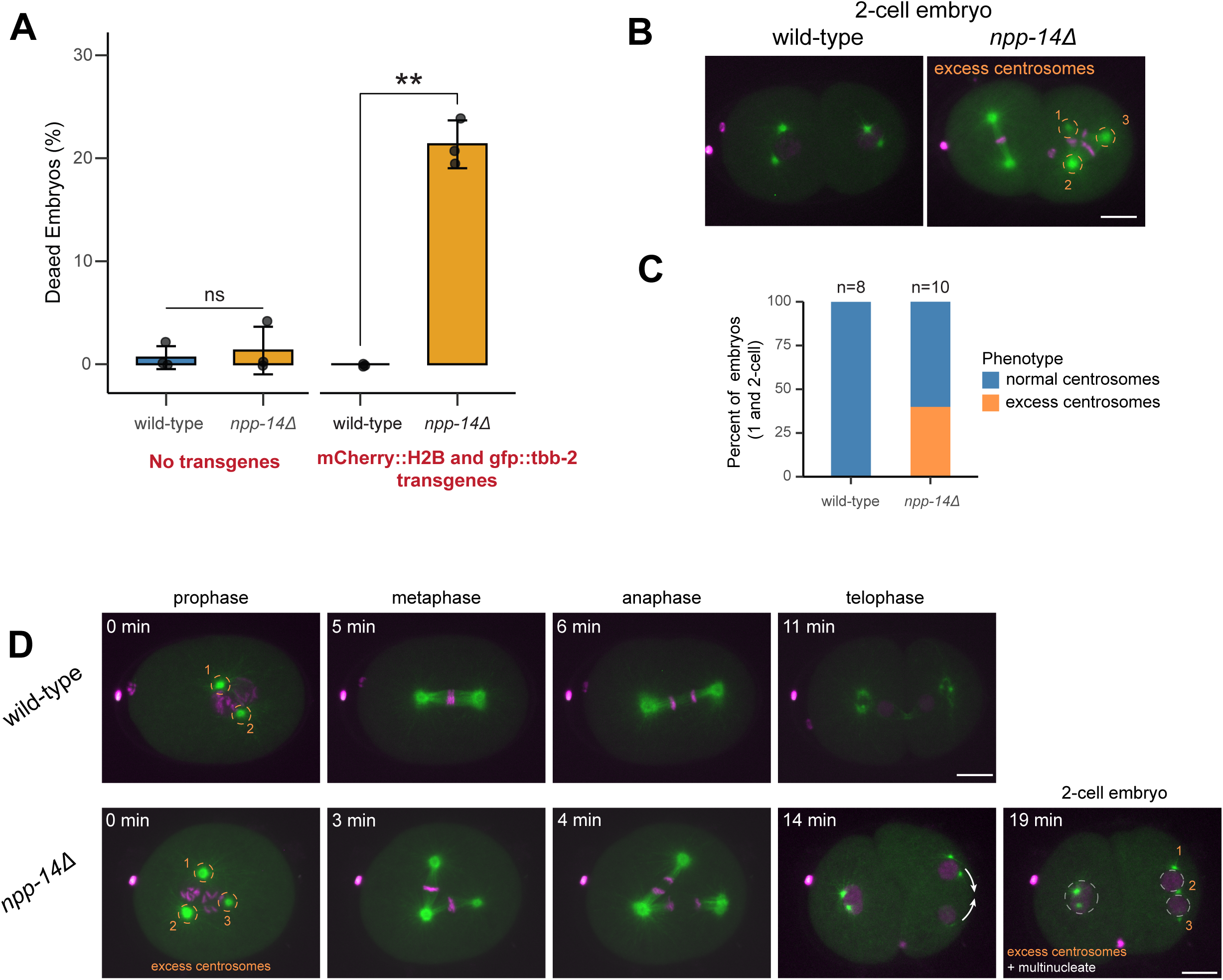
*npp-14* mutants display embryonic defects from excess centrosomes. **A.** Barplot showing mean percentage of dead embryos for wild-type or *npp-14(ax4543)* strains in the N2 background (left) or carrying the *mCherry::h2b*; *gfp::tbb-2* transgene (right) at 20°C. Dots represent biological replicates of approximately 50 embryos, and error bars represent SD. Statistics determined using Student’s t-test. ns = not significant; ** p<0.01. **B.** Maximum intensity projections of 2-cell embryos for the indicated strains showing an extra centrosome in *npp-14(ax4543)*. Dashed orange circles indicate three centrosomes found in a single cell. Scale bar: 10 µm. **C.** Stacked barplot showing percent of embryos with excess centrosomes as shown in B for *npp-14(ax4543)* (n=10) compared to wild-type (n=8) 1 and 2-cell embryos. **D.** Maximum intensity projection timelapse (60x objective) of the first cell division of wild-type vs. *npp-14(ax4543)* embryos expressing mCherry::H2B and gfp::TBB-2. Dashed orange circles indicate centrosomes. In *npp-14(ax4543*), excess centrosomes result in multipolar spindles at metaphase. This leads to two daughter nuclei in a single cell during telophase which come together (as indicated by the solid white arrows), resulting in a multinucleate two-cell embryo. For the 2-cell embryo, white dashed circles indicate nuclei with excess centrosomes numerated in orange. Scale bar: 10 µm.

