## Supplemental Table 1 for "Kinetochore components are perinuclear and form condensates in *C. elegans* germline"

**Table S1. *C. elegans* Strains**

| Name | Genotype | Additional Information | Source |
|---|---|---|---|
| | N2 | wild type | Caenorhabditis Genetics Center (CGC) |
| WHY10 | glh-1(how3[3xHA::TurboID::glh-1]) I | | Price et al., 2021 |
| WHY742 | npp-11(ax4547[npp-11::wrmScarlet]) I; hcp-1(syb441[GFP::hcp-1]) V | npp-11::wrmScarlet from Thomas *et al*., 2023; GFP::HCP-1 from *Edwards et al., 2018* | this study |
| WHY744 | ran-2(ax4545[ran-2::wrmScarlet]) III; hcp-1(syb441[GFP::hcp-1]) V | ran-2::wrmScarlet from Thomas *et al.*, 2023 | this study |
| WHY771 | hcp-1(syb441[GFP::HCP-1]) V | | this study |
| WHY773 | npp-14(ax4543) I; hcp-1(syb441[GFP::HCP-1]) V | npp-14(ax4543) from Thomas *et al*., 2023 | this study |
| WHY889 | npp-14(ax4543) I; ltIs37[pie-1p::mCherry::his-58 + unc-119(+)]; ojIs1[pie-1p::GFP::tbb-2 + unc-119(+)] | | this study |
| WHY891 | ltIs37[pie-1p::mCherry::his-58 + unc-119(+)]; ojIs1[pie-1p::GFP::tbb-2 + unc-119(+)] | | Connolly et al., 2015 |
| WHY910 | npp-14(ax4543) I | | Thomas et al., 2023 |
| WHY1027 | pgl-1(gg547[pgl-1::3xflag::tagRFP]) IV; hcp-1(syb441[GFP::hcp-1]) V | pgl-1::3xflag::tagRFP from Wan *et al.,* 2018 | this study |
| WHY1239 | npp-11(ax4547[npp-11::wrmScarlet]) I; npp-10(ax4538[mNeonGreen::npp-10]) | mNeonGreen::npp-11 from Thomas *et al.*, 2023, how54 deletes the whole ORF and was generated by CRISPR | this study |
| WHY1241 | npp-14(how55) I; npp-11(ax4547[npp-11::wrmScarlet]) I; hcp-1(syb441[GFP::hc | how55 deletes the whole ORF and was generated by CRISPR | this study |
| WHY1273 | 1(syb441[GFP::hcp-1]) V | npp-14(ax4541) from Thomas *et al*., 2023 | this study |
| WHY1281 | npp-11(ax4547[npp-11::wrmScarlet]) I; hcp-1(how56[GFP::HCP-1(Δbipartite nls | how56 deletion of predicted bipartite nls (aa 18-41) by CRISPR | this study |
