## Supplemental Table 2 for "Kinetochore components are perinuclear and form condensates in *C. elegans* germline"

**Table S2: List of oligonucleotides**

| ID | Sequence (5' to 3') | Description |
|---|---|---|
| WT2824 | AATCGACGTTTCTTTCACAA | 5' guide RNA *hcp-1* N-terminus |
| WT2825 | TTCATGGCACATATGTTAGT | 3' guide RNA *hcp-1* C-terminus |
| WT3286 | taatttatagataaatcgacgtttctttcacaATG**AAGCTT**TAGgctataagattctctcaccccactagatctct | ssdonor *hcp-1(how54)* , delete whole ORF + HINDIII restriction site |
| WT3544 | TTGAGGAGAAAATGTTCAAA | 5' guide RNA *npp-14* N-terminus |
| WT2729 | AACCGACATCTTTCACATCA | 3' guide RNA *npp-14* C-terminus |
| WT3576 | AAAATGGCGACTCGGTTTGC | guide RNA *hcp-1* exon 2, antisense, for deleting predicted NLS |
| WT3577 | tttaattaaatcaatgacaatataaattatttcagATTTTTGGAACTTCGTCGGGCAATATGGCCGATAG | ssdonor *hcp-1* (Δpredicted NLS, aa 18-41), paired with guide WT3576 |
| WT1333 | ttgttgccacataacgtcaaa | genotyping sense *hcp-1* upstream ORF |
| WT2835 | tgcgaaaggagagaacatcga | genotyping antisense *hcp-1* downstream ORF |
| WT3617 | GACAACGAGAACAAGCGCTC | genotyping sense *hcp-1* exon 1 |
| WT1334 | GTTGGTTCCCACGAAGGAT | genotyping antisense *hcp-1* exon 2 |
| WT2376 | TGCAGAAGCTGATGGATCTC | genotyping sense *npp-11* exon 5 |
| WT2377 | tggtgatcatgaagaaaacagg | genotyping antisense *npp-11* 3' UTR |
| WT1273 | TCAGTGGATTTGGGGAAGCC | genotyping sense *pgl-1* exon 8 |
| WT1274 | atcattcggcaccgtcgatt | genotyping antisense *pgl-1* downstream ORF |
| WT2370 | tttcgcaatttcgctaacct | genotyping sense *npp-14* upstream ORF |
| WT2372 | atgaagttgctttccgttcg | genotyping antisense *npp-14* downstream ORF |
| WT2374 | TGAAACTGATTGGGGATATGG | genotyping sense *ran-2* exon 10 |
| WT2375 | aaatcgcgaagactgggata | genotyping antisense *ran-2* 3' UTR |
